# Nox4 Mediates Diastolic Function in a Genetic Model of Pitx2 Haploinsufficiency

**DOI:** 10.64898/2026.06.30.735639

**Authors:** Sumner Gardner, Anam Fatima, Faris Abusharkh, Elizabeth Kobeck, Chitra Basu, Francis J. Miller, Vineet Agrawal

## Abstract

Heart failure with preserved ejection fraction (HFpEF) commonly coexists with atrial fibrillation (AF), but shared mechanisms remain unclear. In this study, we hypothesized that *Pitx2*, a transcription factor located near the strongest genetic locus associated with AF in humans, increases susceptibility to HFpEF-like remodeling. We also sought to understand pathways that might be central to this increased risk. Male and female *Pitx2*+/− mice and wild-type littermates received 3-week subcutaneous osmotic pump infusion of saline or angiotensin II (Ang II; 500 ng/kg/min). Cardiac structure and function were assessed by echocardiography and catheterization, and functional capacity by exercise treadmill. RNA transcriptomic profiling was performed to identify candidate pathways. In a separate cohort, Ang II–treated mice were randomized to oral GKT136901 (30 mg/kg/day) or vehicle during infusion. After Ang II infusion, *Pitx2*+/− mice developed exaggerated HFpEF-like changes, including greater left ventricular hypertrophy, left atrial enlargement, diastolic dysfunction, elevated left ventricular end-diastolic pressure, and reduced treadmill performance. RNA-seq showed enrichment of metabolic and stress-response pathways with selective upregulation of *Nox4*, confirmed by RT-qPCR. GKT136901 attenuated structural remodeling, diastolic dysfunction indices, elevated filling pressures, and cardiomyocyte hypertrophy, but did not improve endurance. These findings implicate redox signaling, including *Nox4*, in AF genetic susceptibility–HFpEF interactions.

## INTRODUCTION

Heart failure with preserved ejection fraction (HFpEF) now accounts for 50% of hospitalized heart failure patients and is associated with similar morbidity and mortality to the other forms of heart failure, yet currently has no therapies that significantly improve mortality.(Shah et al., 2017) Among the variety comorbid conditions that are associated with HFpEF, atrial fibrillation (AF) and HFpEF are increasingly recognized as intertwined cardiovascular syndromes that frequently coexist. AF is present in nearly half of all HFpEF patients, and unlike other forms of heart failure, associated with a 40% greater risk of recurrent hospitalization or mortality over 2 years.(Zafrir et al., 2018, Sartipy et al., 2017) Conversely, incident AF is a selective and independent risk for development of HFpEF but not other forms of heart failure.(Santhanakrishnan et al., 2016, Patel et al., 2021)

While strong epidemiologic associations suggest potential shared pathophysiologic mechanisms between AF and HFpEF, the specific shared pathophysiologic mechanisms are not well understood. Genetic studies have identified numerous loci associated with AF susceptibility, among which paired-like homeodomain transcription factor 2 (*PITX2*) represents the strongest and most consistently replicated signal.(Roselli et al., 2025) While the role of *PITX2* in arrhythmogenesis has been extensively studied, recent evidence also suggests that the effects of *PITX2* extend beyond arrhythmias alone. *PITX2* plays an important role in heart development as well as in the mediation of calcium handling, oxidative stress, and myocardial metabolism post-natally.(Hill et al., 2019, Li et al., 2018, Tao et al., 2016, Subati et al., 2025)

Murine models with Pitx2 insufficiency exhibit changes in mitochondrial function, reactive oxygen species generation, and metabolic pathways that resemble molecular signatures observed in heart failure.(Subati et al., 2025, Agrawal et al., 2023) How inherited risk for AF alone or in combination with established common risk factors accelerates myocardial remodeling to lead to HFpEF is not well understood. Thus, the goal of this study was to leverage a transgenic mouse model of *PITX2* haploinsufficiency with and without hypertensive stress, the most common comorbidity associated with HFpEF,(Tsioufis et al., 2017) to gain insight into biologic mechanisms that might drive HFpEF risk in the setting of genetic susceptibility to AF.

## METHODS

### Study Approval

All animal studies were conducted in accordance with and approved by the Vanderbilt University Institutional Animal Care and Use Committee (Vanderbilt University Medical Center IACUC Protocol Number M2200044, Nashville VA Medical Center IACUC Protocol Number V2200053).

### Angiotensin II Pump Infusion

All mice in the current study were bred on the C57BL6/J background. Equal numbers of male and female mice (n=5 each, n=10 per genotype) heterozygous for a *Pitx2*-null allele that removes all isoform function(Wang et al., 2010, Ai et al., 2006) (*Pitx2^+/−^)* were studied at 8 to 10 weeks of age, with age-matched wild-type littermates (*Pitx2^+/+^)* serving as controls. Surface ECG was assessed at the time of echocardiography for all mice and noted to be in sinus rhythm for all mice. At 8-10 weeks of age, mice of both genotypes were randomized to undergo subcutaneous implantation of osmotic pumps (Alzet LLC, Campbell, CA, USA) to deliver saline carrier control or Angiotensin II (HY-13948, MedChemExpress, Monmouth Junction, NJ, USA) at 500 ng/kg/min for 3 weeks. Mice underwent sedated echocardiography at baseline and prior to the endpoint of the study. They additionally underwent treadmill exercise testing in the final week prior to the endpoint of the study. At sacrifice, all mice underwent non-survival open chested left and right heart catheterization for invasive hemodynamic assessment.

### Mouse Treadmill Endurance Testing

All mice underwent treadmill endurance testing on a variable speed 6-chamber treadmill (Columbus Instruments, Columbus, OH, USA) in the final week prior to the endpoint of the study. To acclimate the mice to the treadmill, they underwent two consecutive days of exposure to the treadmill at a low speed of 10 m/min for 10 minutes per day. On the third day, mice underwent a stress test protocol with incremental increase in treadmill speed by 1 m/min every 2 minutes to define maximal tolerated speed per mouse. On the fourth day, the mice then underwent an endurance test where treadmill speed started at 8 m/min x 10 minutes, followed by 2 m/min increase every 5 minutes until 60% of the maximal speed was obtained. Thereafter, the mice were allowed to continue to walk until exhaustion, and total exercise distance was calculated.

### Mouse Echocardiography and Catheterization

Mouse echocardiography and catheterization was performed as previously reported.(Agrawal et al., 2019) For echocardiography, studies were performed at baseline prior to subcutaneous pump implantation and one day prior to sacrifice. For echocardiography, mice were anesthetized with isoflurane 2-3% and depilatory cream was applied to the thorax. Mice were then placed on a heated table in supine position and images were acquired using the VisualSonics Vevo F2 Platform in B-mode, M-mode, and Doppler mode. Parasternal long-axis, short-axis, and modified RV-centric views were obtained as previously described.(Brittain et al., 2013) Thereafter, mice were allowed to recover for 24 hours before undergoing anesthesia again with 2-3% isoflurane for open chest catheterization. Mice were orotracheally intubated with 22g catheter, mechanically ventilated at 18 cc/kg, and anesthetized with 2-3% vaporized isoflurane general anesthesia. On a heated surgical table, the diaphragm was surgically exposed through a ventral incision in the abdomen. After takedown of the diaphragm, a 1.4 French Mikro-tip catheter was directly inserted into the left ventricle for measurement of pressure and volume within the left ventricle, followed by the right ventricle. Hemodynamics were continuously recorded with a Millar MPVS-300 unit coupled to a Powerlab 8-SP analog-to-digital converter acquired at 1000 Hz and captured to a Macintosh G4 (Millar Instruments, Houston, TX). Left ventricular tissue was then harvested for downstream studies.

### RNA sequencing

Left ventricular tissue was harvested from mouse studies and immediately placed into RLT buffer (Qiagen, Hilden, Germany). RNA isolation was then performed using the Qiagen RNeasy Plus Mini Kit (Qiagen, Hilden, Germany) following manufacturer’s guidelines and delivered to Novogene (Sacramento, California, USA) for paired-end sequencing on an Illumina platform. A nominal read depth of 20 million RNA (40 million ends) per sample were used. Initial alignment and quantification were performed using the Partek Flow package. Gene ontology analysis was conducted using Webgestalt.

### Nox1/4 Inhibition

Equal numbers of male and female mice *Pitx2^+/−^* or *Pitx2^+/+^* littermate controls underwent subcutaneous Ang-II pump implantation as described above for 3 weeks. Thereafter, mice were randomized to daily oral gavage of Nox1/4 inhibitor, GKT136901 (HY-101499, MedChemExpress, Monmouth Junction, NJ, USA) administered to 30 mg/kg or carrier control for the duration of the 3-week study. GKT136891 was solubilized to a 30 mg/ml solution in DMSO, and mixed 1:9 with corn oil to a working dilution of 3 mg/ml.(Sedeek et al., 2013) Mice underwent baseline echocardiography, and they underwent exercise treadmill testing, echocardiography, and catheterization at the endpoint of the study.

### Immunostaining

Hearts were perfused with 50 mM KCl in 5% dextrose to fix hearts in diastole, removed, and immediately fixed in 4% paraformaldehyde for 10 minutes prior to transferring to 30% sucrose for 24 hours. Afterwards, hearts were cut in transverse sections and OCT embedded for frozen sectioning (5 *μ*m) and wheat germ agglutinin staining. Wheat germ agglutinin staining was performed by permeabilizing sections (1% BSA, 0.3% TritonX, 5% goat serum) for 30 minutes, incubating slides in 100 *μ*g/ml of Alexa Fluor 488-conjugated wheat germ agglutinin (Invitrogen, Carlsbad, CA, USA; W11261) for 1 hour in the dark, and then counterstaining with Prolong Gold Antifade with DAPI (Invitrogen, P36931) before curing for 24 hours. Imaging was performed on a Keyence BZ-X800 microscope. Quantification of cross-sectional area on WGA-stained slides was performed in a blinded fashion using the Fiji distribution of ImageJ.(Schindelin et al., 2012)

### Statistical Analysis

All statistical analysis was conducted using one-way or two-way ANOVA in GraphPad Prism 10.

### Data Availability

RNA sequencing data will be deposited to the NCBI GEO database. All other data is available upon reasonable request to the corresponding author.

## RESULTS

### Pitx2 +/− Mice Develop Features of Heart Failure with Preserved Ejection Fraction with Ang II Infusion

We first sought to characterize the baseline cardiac structure and function of *Pitx2^+/−^* and *Pitx2^+/+^*mice. At baseline, we identified no differences in cardiac structure or function between haploinsufficient or wild type littermate controls (**Figure 1**), consistent with prior studies.(Subati et al., 2025) With Ang-II infusion for 3 weeks, we found that all mice demonstrated a normal left ventricular ejection fraction (**Figure 1A**). However, with Ang II infusion but not at baseline, Pitx2^+/−^ mice displayed decreased cardiac output (**Figure 1B**), decreased end-diastolic volume of the left ventricle (**Figure 1C**), increased left ventricular posterior wall thickness (**Figure 1D**), increased left atrial end-systolic volume (**Figure 1E**), and abnormally increased E/e’ ratio (**Figure 1F**).

**Figure 1.**
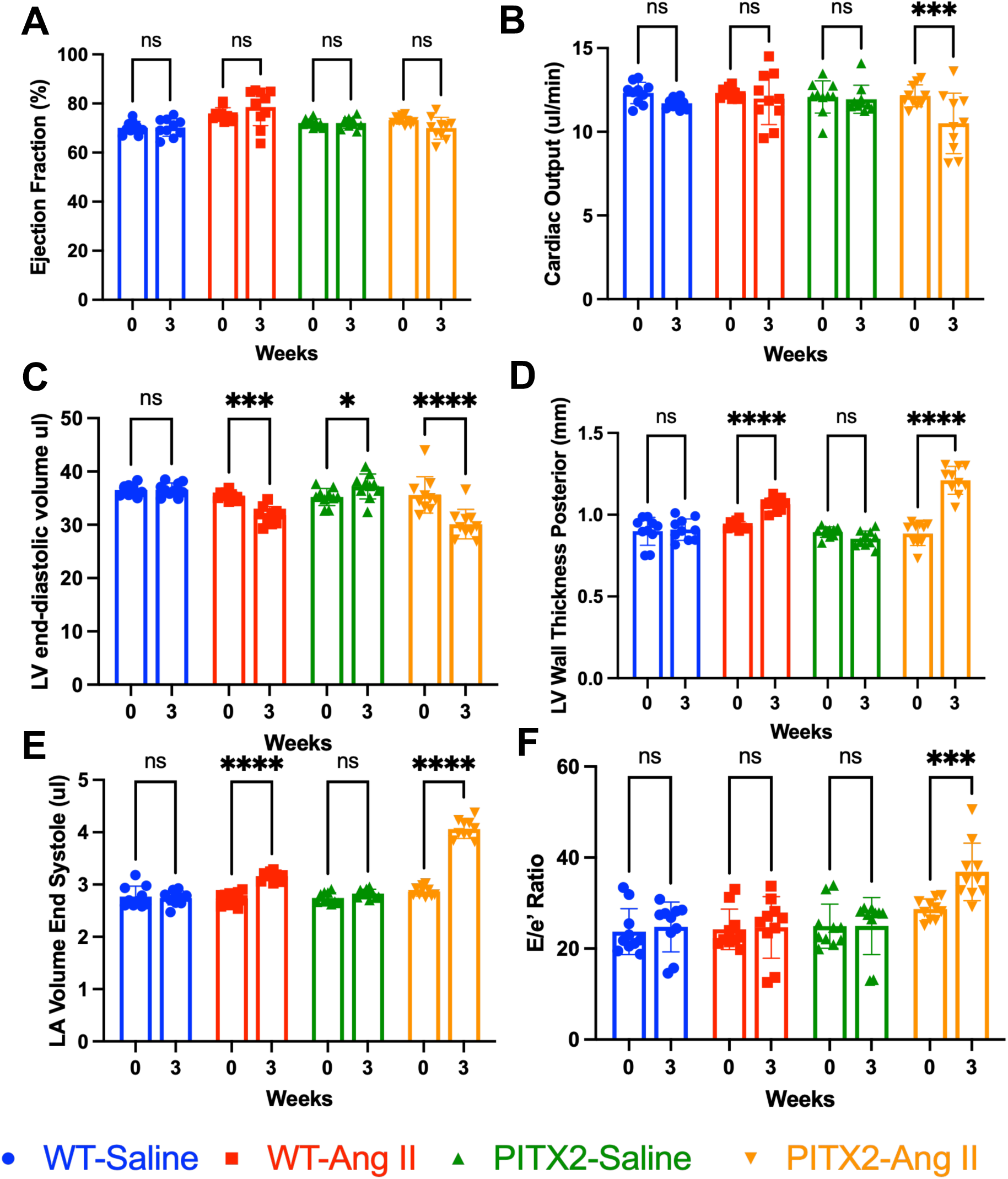
Echocardiographic measure of (A) left ventricular ejection fraction, (B) cardiac output, (C) left ventricular end-diastolic volume, (D) left ventricular posterior wall thickness, (E) left atrial end-systolic volume, and (F) E/e’ ratio in either Pitx2 haploinsufficient or wildtype littermate control mice subjected to Angiotensin II infusion by osmotic pump or saline carrier control for 3 weeks. *LV = left ventricle, LA = left atrium. * p < 0.05, ** p < 0.01, *** p < 0.001, **** p < 0.0001*.

We next sought to further characterize the cardiac structural changes with invasive hemodynamics. Ang II had a reproducible effect of increased systolic blood pressure and no effect on heart rate at the time of catheterization, irrespective of genotype (**Figure 2A,B**). When compared to littermate controls, *Pitx2^+/−^* disproportionately increased left ventricular end-diastolic pressure and left ventricular relaxation constant (**Figure 2 C,D**). Ang II treatment also increased cardiomyocyte cross-sectional area in all mice, but to a greater extent in *Pitx2^+/−^* mice as compared to their wild-type littermate controls (**Figure 2E**). *Pitx2^+/−^* mice, as compared to their wild type counterparts, also displayed diminished exercise capacity when exposed to Ang II infusion (**Figure 3**).

**Figure 2.**
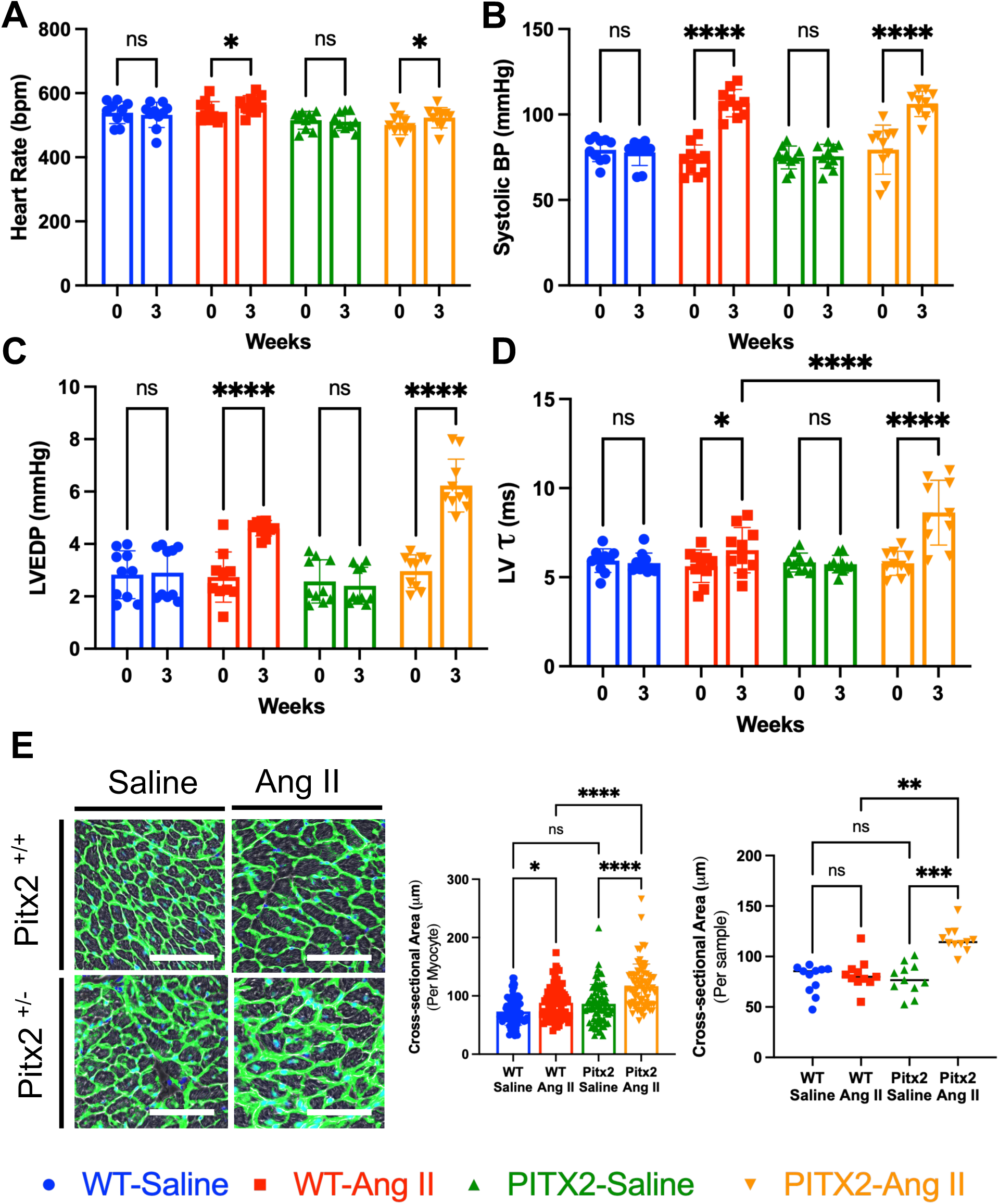
Invasive catheterization-based measurement of (A) heart rate, (B) systolic blood pressure (estimated by left ventricular systolic pressure and confirmed by aortic tracing), (C) left ventricular end-diastolic pressure, (D) left ventricular diastolic relaxation constant (*τ*), and (E) cardiomyocyte cross-sectional area measured per individual myocytes as well as per sample in Pitx2 haploinsufficient or wildtype littermate control mice subjected to Angiotensin II infusion by osmotic pump or saline carrier control for 3 weeks. *BP = blood pressure, LVEDP = left ventricular end-diastolic pressure. * p < 0.05, ** p < 0.01, *** p < 0.001, **** p < 0.0001*.

**Figure 3.**
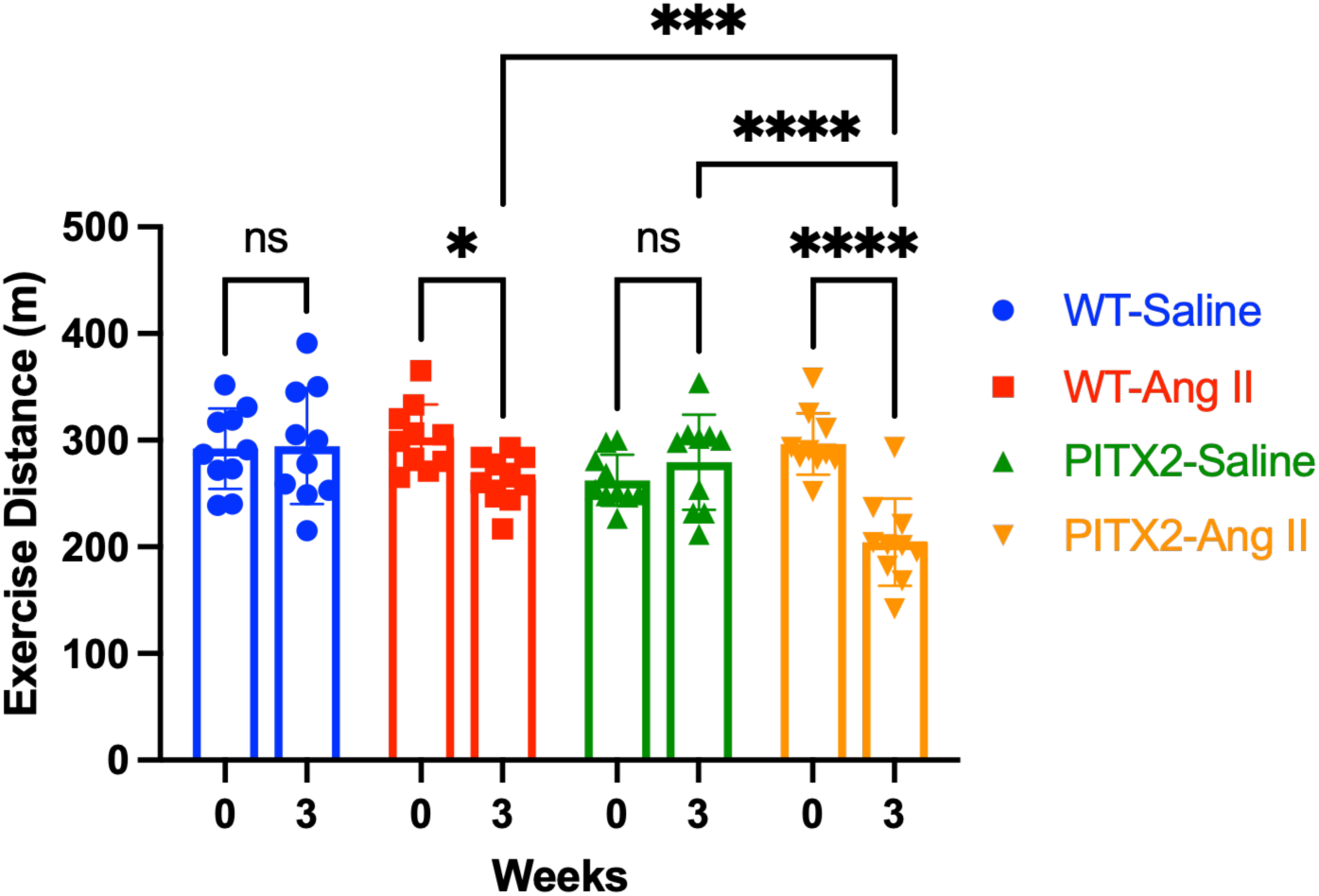
Treadmill-based exercise endurance tolerance, as measured by total distance exercised, in Pitx2 haploinsufficient or wildtype littermate control mice subjected to Angiotensin II infusion by osmotic pump or saline carrier control for 3 weeks. *\* p < 0.05, ** p < 0.01, *** p < 0.001, **** p < 0.0001*.

### Nox4 Is Upregulated in Hearts of Pitx2^+/−^ Mice After Ang II Infusion

We next sought to identify transcripts differentially expressed in *Pitx2^+/−^* vs *Pitx2^+/+^* mice subjected to Ang II infusion. After quality control of the sequenced data, we identified 10,867 unique transcript with at least 1 sequenced transcript per million. At an unadjusted p-value threshold of 0.05, we identified 92 genes that were differentially expressed in *Pitx2^+/−^*mice (**Figure 4A**). Gene set enrichment analysis revealed that the most significantly overrepresented gene ontology (GO) terms in the biological process domain were primarily associated with metabolic regulation and homeostasis, with additional GO terms associated with cellular communication, cellular stress response, and regulation of the immune system (**Figure 4B**).

**Figure 4.**
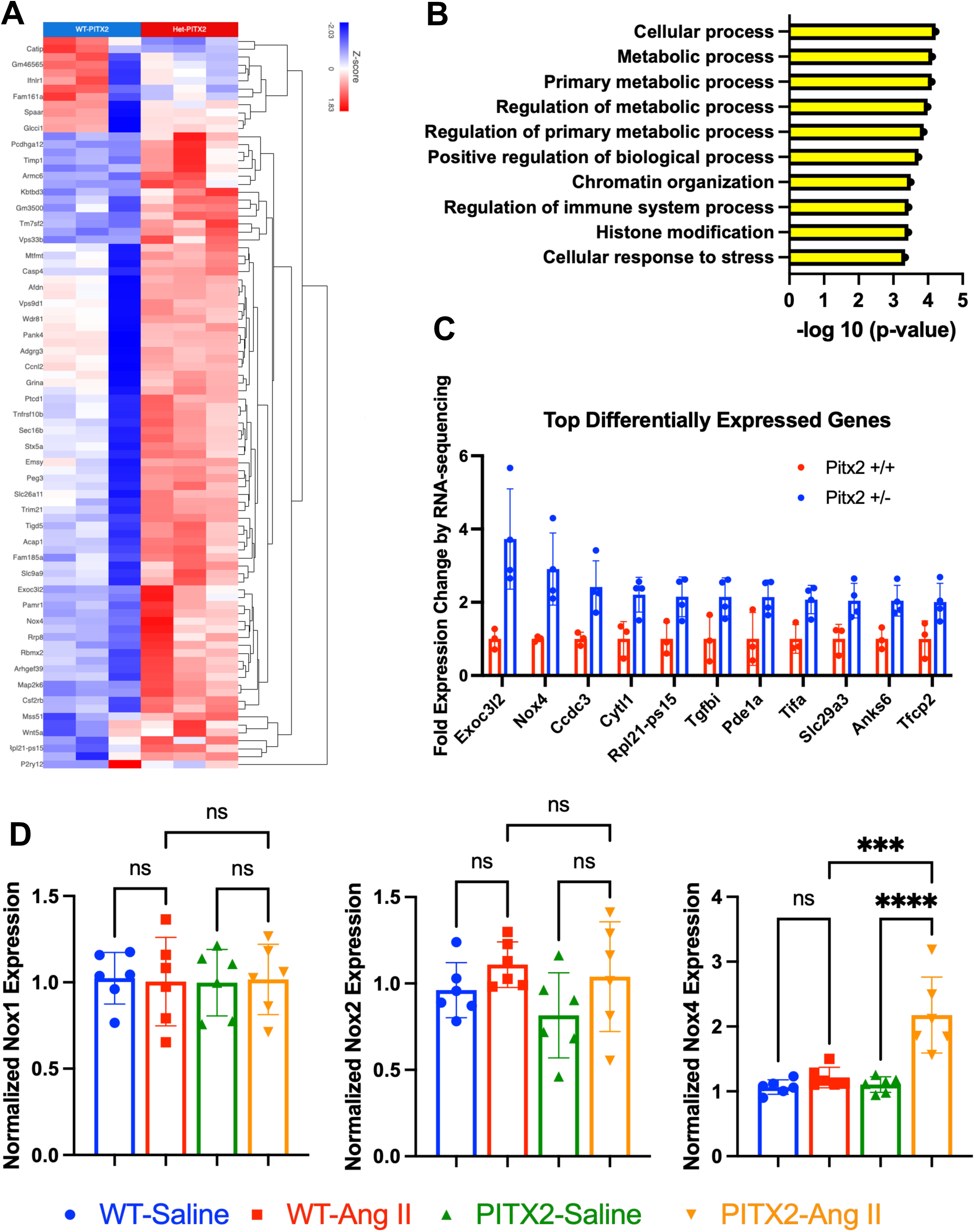
(A) Hierarchical clustering heat map of differentially expressed transcripts in the left ventricles of Pitx2 haploinsufficient or wild type mice subjected to Angiotensin II infusion. (B) Top over-represented gene ontologies in gene set enrichment analysis. (C) Top 11 differentially expressed genes. (D) Reverse transcriptase quantitative polymerase chain reaction of NADPH oxidase enzyme isoforms to validate transcriptional changes identified by RNA-sequencing. *\*\*\* p < 0.001, **** p < 0.0001*.

Among the top 92 differentially expressed genes, only 11 genes displayed at least a 50% change in expression. Among those 11 differentially expressed genes, the gene encoding *Nox4* was one of the top differentially expressed genes in *Pitx2^+/−^* mice (**Figure 4C**). Given the known associations between *Pitx2* isoforms and regulation of oxidative stress-related pathways,(Sridhar et al., 2024, Steimle et al., 2022) we verified findings from RNA-sequencing with RT-qPCR of left ventricular tissue and found that the only isoform of *Nox* genes that was differentially expressed in *Pitx2^+/−^* mice subjected to Ang II infusion was *Nox4*, with no significant changes among other transcripts (**Figure 4D**).

### Nox1/4 Inhibition Mitigates Cardiac Remodeling in Pitx2^+/−^ Mice After Ang II Infusion

To understand the contribution of *Nox4* to cardiac remodeling, we randomized *Pitx2^+/−^* or *Pitx2^+/+^* mice to Ang II infusion with or without oral administration of GKT136901, a *Nox1/4* inhibitor,(Sedeek et al., 2013) for 3 weeks. After 3 weeks of treatment, all mice demonstrated normal ejection fraction and no significant differences in cardiac output (**Figure 5A,B**). *Pitx2^+/−^*mice treated with GKT136901 did, however, show an improvement in left ventricular end-diastolic volume, left ventricular posterior wall thickness, reduction of left atrial end-systolic volume, and improvement in E/e’ ratio (**Figure 5C-E**). By invasive catheterization, GKT136901 had no effect on heart rate or systolic blood pressure (**Figure 6A,B**), but did improve left ventricular end-diastolic pressure, left ventricular diastolic relaxation, and cardiomyocyte hypertrophy as measured by cross-sectional area (**Figure 6C-E**). Notably, though, GKT1136901 did not improve exercise capacity in *Pitx2^+/−^* mice compared to wild-type mice (**Figure 7**).

**Figure 5.**
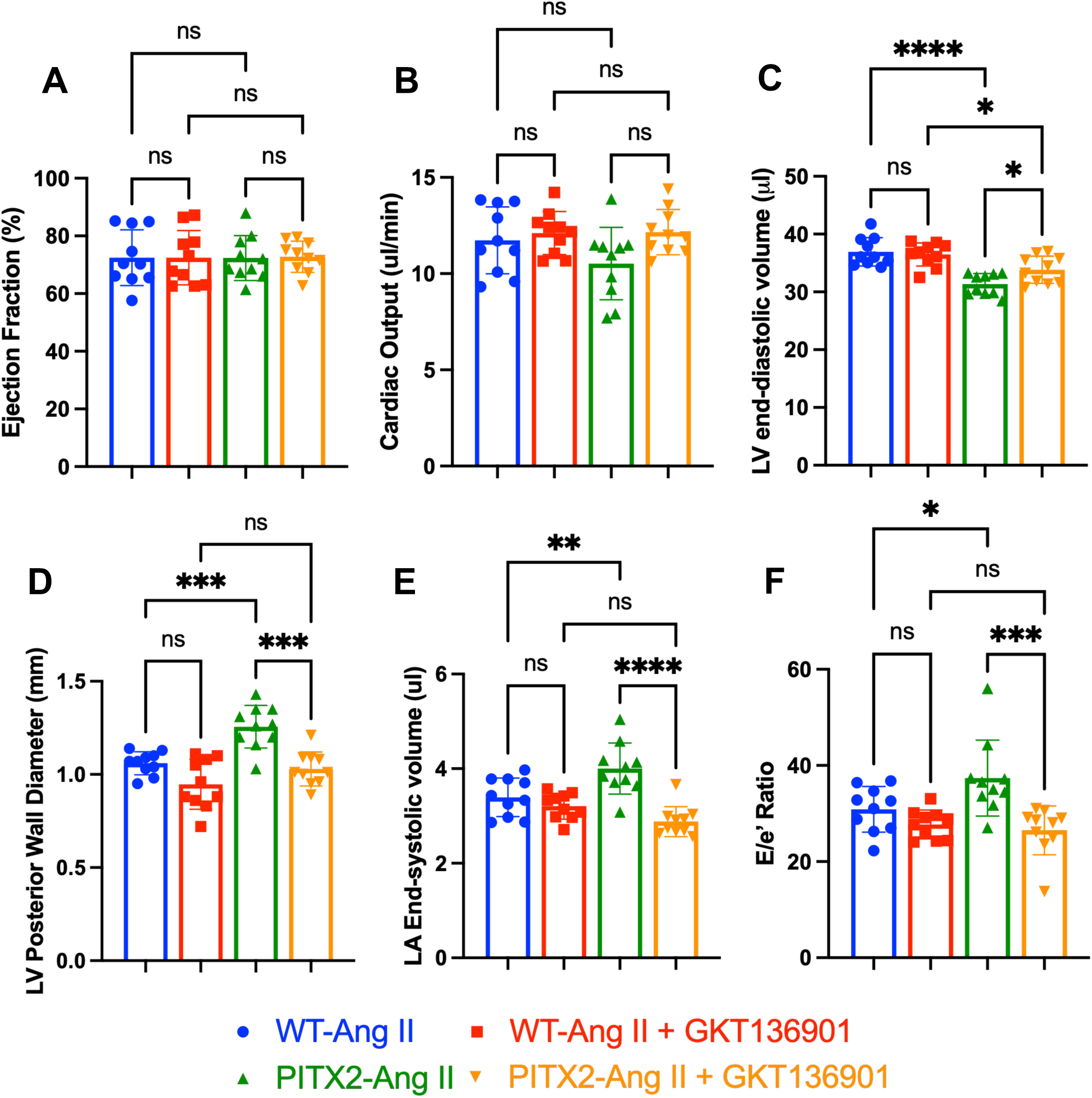
Echocardiographic measure of (A) left ventricular ejection fraction, (B) cardiac output, (C) left ventricular end-diastolic volume, (D) left ventricular posterior wall thickness, (E) left atrial end-systolic volume, and (F) E/e’ ratio in either Pitx2 haploinsufficient or wildtype littermate control mice subjected to Angiotensin II infusion and treatment with either GKT136901 or carrier control for 3 weeks. *LV = left ventricle, LA = left atrium. * p < 0.05, ** p < 0.01, *** p < 0.001, **** p < 0.0001*.

**Figure 6.**
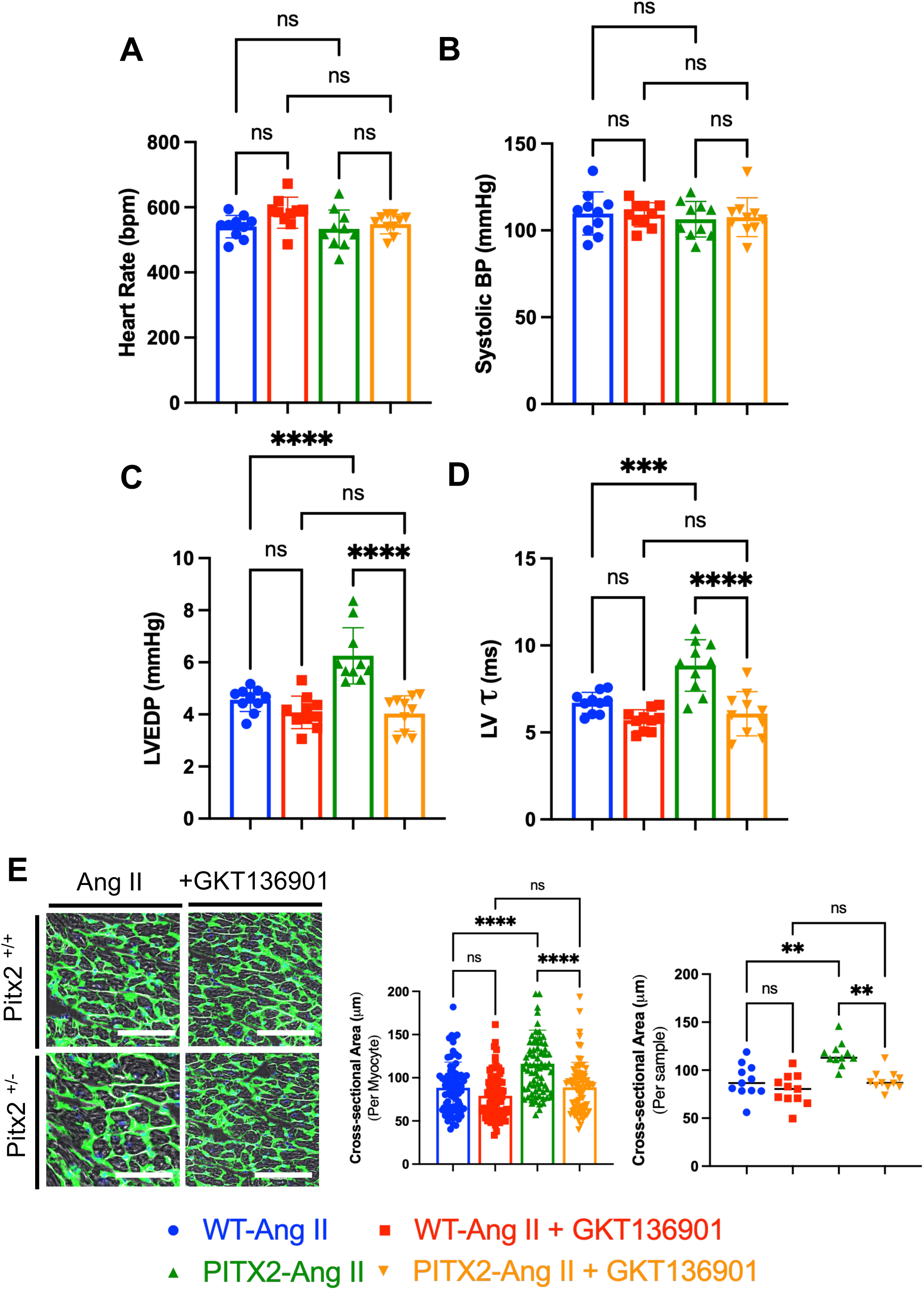
Invasive catheterization-based measurement of (A) heart rate, (B) systolic blood pressure (estimated by left ventricular systolic pressure and confirmed by aortic tracing), (C) left ventricular end-diastolic pressure, (D) left ventricular diastolic relaxation constant (*τ*), and (E) cardiomyocyte cross-sectional area measured per individual myocytes as well as per sample in Pitx2 haploinsufficient or wildtype littermate control mice subjected to Angiotensin II infusion and treatment with either GKT136901 or carrier control for 3 weeks. *BP = blood pressure, LVEDP = left ventricular end-diastolic pressure. * p < 0.05, ** p < 0.01, *** p < 0.001, **** p < 0.0001*.

**Figure 7.**
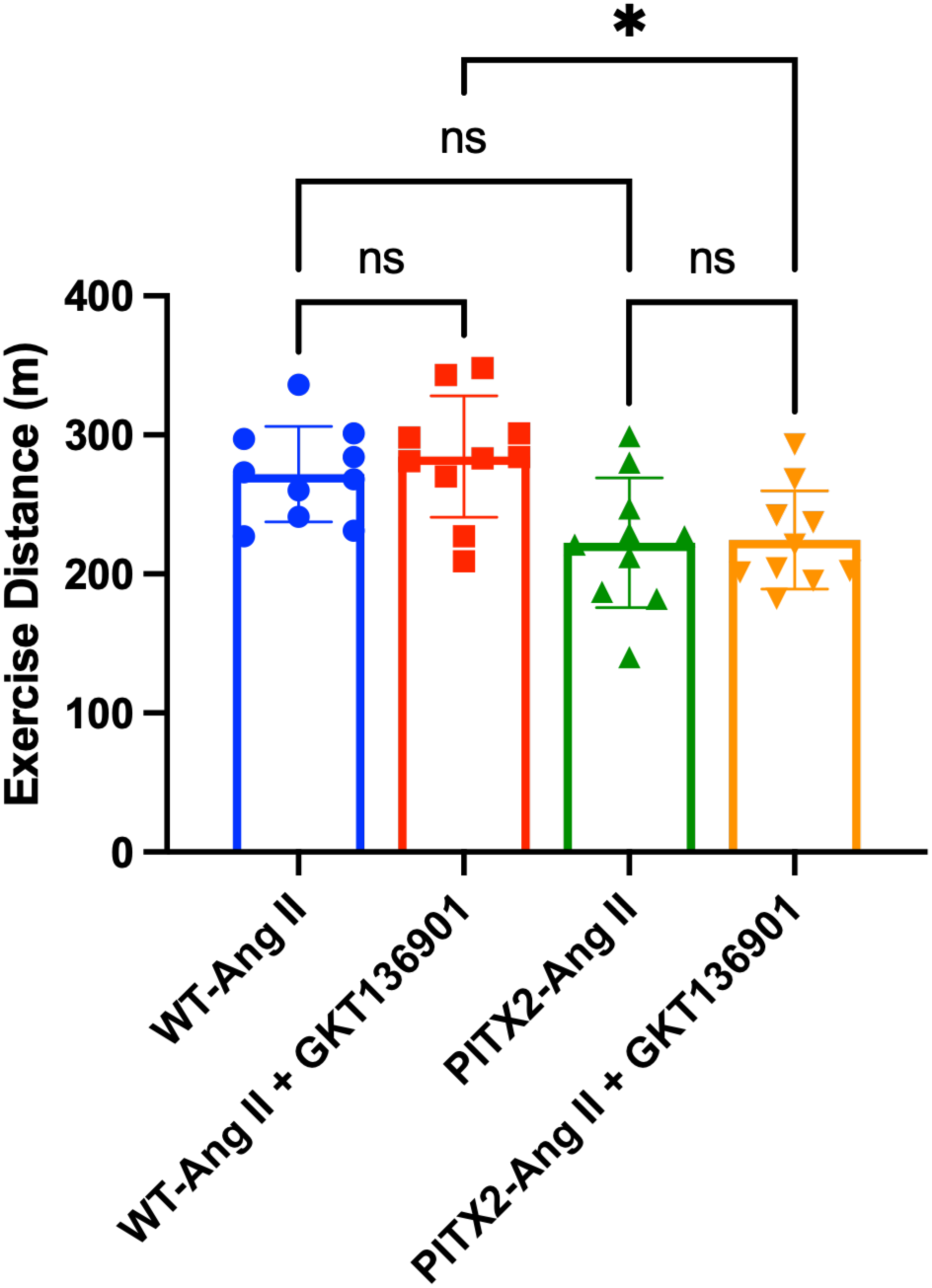
Treadmill-based exercise endurance tolerance, as measured by total distance exercised, in Pitx2 haploinsufficient or wildtype littermate control mice subjected to Angiotensin II infusion and treatment with either GKT136901 or carrier control for 3 weeks. *\* p < 0.05, ** p < 0.01, *** p < 0.001, **** p < 0.0001*.

## DISCUSSION

In this study, we leveraged a genetic model of atrial fibrillation (AF) susceptibility, Pitx2 haploinsufficiency, combined with a hypertensive stress, the most common comorbidity associated with heart failure with preserved ejection fraction (HFpEF), to understand shared mechanisms that may link AF and HFpEF pathophysiology.(Deichl et al., 2022) Our study has three principal findings. The first is that *Pitx2*^+/−^ mice exposed to Ang II infusion to induce hypertension developed accelerated diastolic dysfunction relative to their wild type littermate counterparts. The second is that transcriptomic profiling of left ventricular tissue identified enrichment of pathways related to metabolic homeostasis and stress responses, and highlighted NOX4 as a prominent differentially expressed transcript in Ang II–treated *Pitx2*^+/−^ mice, with isoform-specific upregulation confirmed by RT-qPCR. Third, GKT136901, a pharmacologic *Nox1/4* inhibitor,(Sedeek et al., 2013) mitigated multiple structural and hemodynamic features of maladaptive remodeling, including LV hypertrophy, LA enlargement, diastolic dysfunction indices, and cardiomyocyte hypertrophy, although it did not improve treadmill exercise capacity over the study timeframe.

Reactive oxygen species comprise a family of unstable oxygen-containing molecules that are not only produced within cells as a byproduct metabolism, but also serve as important molecules within the heart to regulate excitation-contraction coupling, calcium handling, mitochondrial function, vascular tones, and adaptive stress responses.(Csanyi and Miller, 2014) NADPH oxidase (NOX) enzymes are a major enzymatic source of these reactive oxygen species (ROS) in the heart, generating superoxide or hydrogen peroxide as signaling molecules that modulate cardiomyocyte and vascular function.(Streeter et al., 2013) Among the five NOX enzyme isoforms found in humans, all but Nox5 are present in rodent models.(Fulton, 2009) Isoforms NOX1, NOX2, and NOX4 are known to be expressed within the heart, with NOX4 being the only enzyme that is constitutively active within cardiomyocytes. NOX4 is expressed primarily on the intracellular membranes of cardiomyocytes, including the endoplasmic reticulum, T-tubule, mitochondria, and perinuclear envelope, where it regulates redox sensitive reactions through generation of hydrogen peroxide.(Gray et al., 2019) While some studies suggest that transient activation of NOX4 is a stress-adaptive signaling in response to stressors,(Zhang et al., 2010, Zhao et al., 2015) other studies have demonstrated that constitutive activation of NOX4 leads to excessive oxidative stress in the multiple compartments where NOX4 is expressed.(Lozhkin et al., 2022, Vendrov et al., 2023)

The role of oxidative stress is well documented in atrial fibrillation (AF). One of the strongest genetic loci associated with AF risk is located near the *PITX2* gene, which has been implicated in several studies as playing a role in regulating oxidative stress programs in cardiomyocytes.(Steimle et al., 2022, Subati et al., 2025) Our study implicates an important role for *Nox4* in potentially regulating this oxidative stress program. Consistent with prior studies of cardiac tissue from patients with AF in which NOX4 was the only NOX isoform increased in expression,(Zhang et al., 2010) our study also identified a selective increase in NOX4 expression in *Pitx2*^+/−^ mice infused with Ang II. Also consistent with the consequences of increased NOX4 expression based on prior work, our study observed an over-representation of gene ontologies associated with metabolic function in the heart altered in *Pitx2*^+/−^ mice infused with Ang II compared to wild type littermates.(Lozhkin et al., 2022)

These findings differ from prior literature investigating AF mechanisms in which the NOX2 isoform has been identified to be increased in expression in a *Pitx2*-dependent fashion after diet induced obesity.(Sridhar et al., 2024) While this study had notable differences – a wild type murine model as opposed to *Pitx2* haploinsufficient model, diet-induced obesity as a primary stimulus vs hypertensive stress, and primary outcome of atrial fibrillation vs diastolic dysfunction in our study – our study highlights the important observation that regulation of NOX enzymes in heart disease is highly context and cell-dependent.

HFpEF is widely recognized as a syndrome driven by the interaction of systemic comorbidities (e.g., hypertension, obesity, diabetes, chronic kidney disease) with cardiac remodeling and impaired diastolic reserve.(Agrawal et al., 2019, Agrawal et al., 2023) In that framework, our data suggest that inherited AF susceptibility due to reduced *Pitx2* expression functions as a “sensitizing” substrate that unmasks vulnerability to HFpEF-like remodeling when paired with a modest hypertensive stress. Prior studies have shown that heterozygous loss of *Pitx2* multiple gene networks in the heart including altering expression of cell-identity associated transcription factors such as *Tbx5* and *Tbx20*, alter oxidative stress-related pathways in cardiomyocytes through disruption of *Nrf2* networks, and alter BMP10-mediated signaling and myocyte-endothelial communication.(Steimle et al., 2022) Our work suggests that oxidative stress-related pathways may be an important, and nodal, gene network that regulates the maladaptive response in the heart to additional stressors such as hypertensive stress, but future work is necessary to dissect out the specific contributions of these various gene networks in response to additional stressors.

Our study showed that pharmacologic inhibition of *Nox4* ameliorated cardiac remodeling after modest hypertensive stress but did not alter exercise capacity. This discordance mirrors clinical HFpEF, where functional limitation is multifactorial and may reflect contributions from peripheral oxygen extraction, skeletal muscle abnormalities, chronotropic incompetence, pulmonary vascular reserve, and right heart-pulmonary coupling in addition to LV filling pressures. *Pitx2* is also known to be expressed in the skeletal muscle, lungs, pituitary glands, and brain, among other tissues. Our study did not investigate consequences of *Pitx2* haploinsufficiency in these tissues, but given that neuronal input, neurohormonal pathways, lung function, and skeletal muscle are all important regulators of oxygen utilization and exercise capacity in organisms, it is highly likely that *Pitx2* haploinsufficiency affects exercise capacity through multifactorial mechanisms. Given that the heart’s contribution to oxygen utilization is typically considered a central factor in determining exercise capacity, though, it is also possible that our treadmill endpoint could be insensitive to modest improvements in exercise capacity over the short time frame of the study.

Several limitations should be considered. First, the sample size was modest, and although multiple orthogonal phenotyping modalities supported a consistent signal, larger studies will be important to confirm effect sizes, explore sex-specific responses, and improve precision. Second, Ang II infusion for three weeks models a specific neurohormonal/hypertensive stressor; HFpEF in humans arises from diverse comorbidity constellations. Whether *Pitx2* similarly amplifies remodeling in other HFpEF-relevant stress models (e.g., obesity/metabolic stress, aging, renal dysfunction) remains unknown. Third, while our study implicates an important role for *Nox4* in regulating cardiac remodeling in the context of *Pitx2* haploinsufficiency, the findings of this study do not implicate a direct regulation of Nox4 by *Pitx2*. Dissecting out the specific mechanisms by which *Pitx2* loss might regulate Nox4 expression requires further work. Fourth, our transcriptomic analysis, while functioning as a discovery-based approach to understand dominant pathways dysregulated by Ang II stress in *Pitx2^+/−^*mice, was limited in that it was performed at a single endpoint and does not address cell-type specificity. Fifth, our study used GKT136901, a dual Nox1/4 inhibitor and demonstrated cardiac structural/functional improvement after Ang II infusion. Since our study found no differences in *Nox1* expression across all experimental groups and a selective increase in *Nox4* only, the findings do likely reflect the inhibition of *Nox4* by GKT136901. However, future studies of genetic loss-of-function or more selective pharmacologic approaches will be needed to strengthen the findings of this study. Finally, while study used a mouse model susceptible to AF after pacing induction, the mice remained in sinus rhythm throughout the study. The explicit goal of this study was to investigate the consequences of genetic susceptibility to AF independent of the AF rhythm itself, which is known to cause hemodynamic stress on the cardiovascular system. As such, our study also did not investigate whether Ang II infusion rendered *Pitx2* haploinsufficient mice to greater AF susceptibility but could be addressed in future studies.

In summary, we found that *Pitx2* haploinsufficiency, an established genetic determinant of AF susceptibility, predisposes to HFpEF-like structural and hemodynamic remodeling under sub pressor Ang II stress, and is associated with transcriptional changes implicating metabolic and stress-response pathways including upregulation of *Nox4*. Pharmacologic *Nox1/4* inhibition attenuated maladaptive remodeling and improved diastolic indices, supporting a contributory role for Nox-dependent redox signaling in this gene–environment interaction model.

**Table 1.**
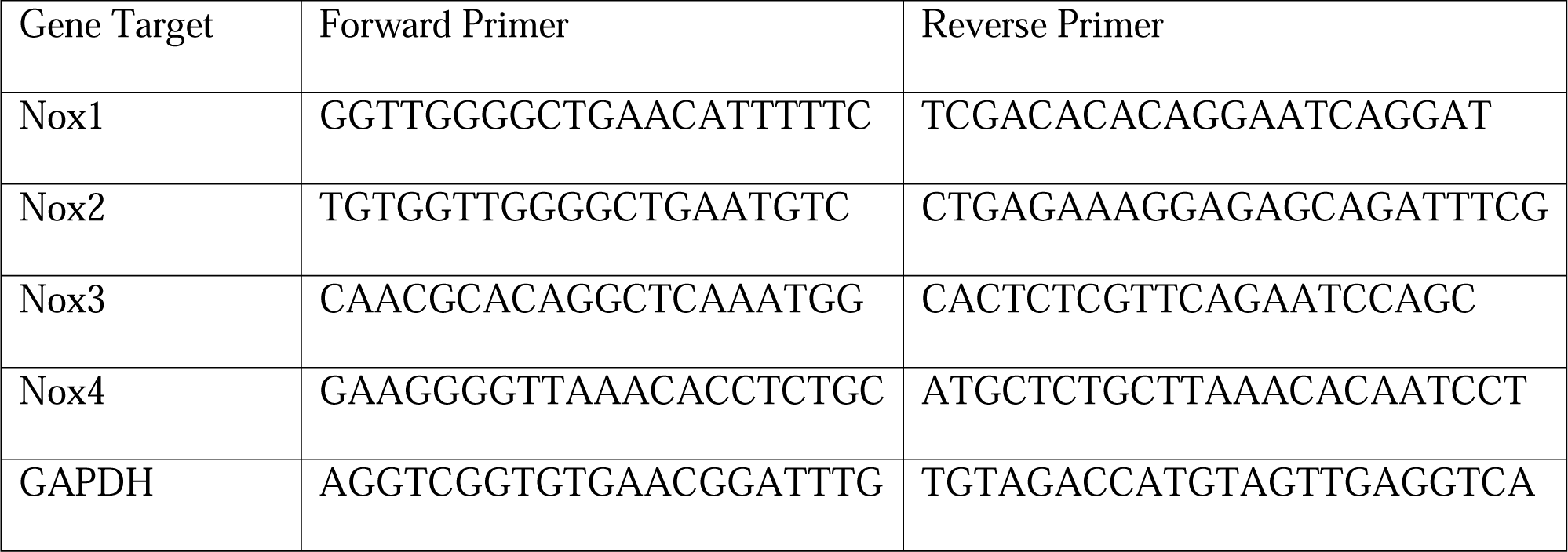
Primer sequences for RT-qPCR studies.

## AUTHOR CONTRIBUTION STATEMENT

SG, AF, FA, and EK all contributed to data curation, formal analysis, investigation, and writing. CB and FJM contributed to formal analysis, methodology, and writing. VA contributed to conceptualization, funding acquisition, data curation, formal analysis, and writing.

## COMPETING INTERESTS STATEMENT

The authors declare there are no competing interests.

## FUNDING STATEMENT

This work was supported by the Department of Veterans Affairs grant 1IK2BX005828 (VA).

## Conflicts of Interest and Disclosures

The contents do not represent the view of the VA or United States Government. The authors otherwise declare no conflicts of interest.

## Notes

### Competing Interest Statement

The authors have declared no competing interest.

